# Microbiota-Independent Antiviral Effects of Antibiotics on Enteric Viruses

**DOI:** 10.1101/2020.02.14.949123

**Authors:** Mikal A. Woods Acevedo, Julie K. Pfeiffer

**Affiliations:** Department of Microbiology, University of Texas Southwestern Medical Center, Dallas, Texas, USA

**Keywords:** coxsackievirus, poliovirus, antibiotics, microbiota

## Abstract

Coxsackieviruses primarily infect the gastrointestinal tract of humans, but they can disseminate systemically and cause severe disease. Using antibiotic treatment regimens to deplete intestinal microbes in mice, several groups have shown that bacteria promote oral infection with a variety of enteric viruses. However, it is unknown whether antibiotics have microbiota-independent antiviral effects for enteric viruses or whether antibiotics influence extra-intestinal, systemic infection. Here, we examined the effects of antibiotics on systemic enteric virus infection by performing intraperitoneal injections of either coxsackievirus B3 (CVB3) or poliovirus followed by quantification of viral titers. We found that antibiotic treatment reduced systemic infection for both viruses. Interestingly, antibiotics reduced CVB3 titers in germ-free mice, suggesting that antibiotic treatment alters CVB3 infection through a microbiota-independent mechanism. Overall, these data provide further evidence that antibiotics can have noncanonical effects on viral infection.

## Introduction

Coxsackievirus B3 (CVB3) is a nonenveloped single-stranded positive-sense RNA virus belonging to the *Enterovirus* genus of the *Picornaviridae* family. CVB3 is spread through the fecal-oral route and has been associated with a wide range of diseases due to systemic viral dissemination, including gastrointestinal distress, type 1 diabetes, myocarditis, aseptic meningitis, cardiac arrhythmias, and death (1-5). Mouse models of CVB3 infection and pathogenesis have used a variety of inoculation routes, including the natural oral route of infection (6, 7), or intraperitoneal (IP) injection, which bypasses the intestinal tract and induces systemic infection of a variety of tissues (8-10).

Recently, we have shown that CVB3 is dependent on a vast intestinal microbial ecosystem of approximately 10^14^ organisms, termed the microbiota, to establish infection in both intestinal and extra-intestinal tissues following oral inoculation (6). Furthermore, we showed that orally delivered CVB3 is relatively sensitive to antibiotic treatment, as a single dose of one antibiotic was sufficient to reduce CVB3 replication and pathogenesis (6). However, it remains unknown if antibiotics reduce systemic CVB3 infection through a microbiota-dependent or independent mechanism.

Our group and others have used antibiotics to deplete the host microbiota to study virus-host-microbiota interactions (6, 11-14). However, antibiotics can have microbiota-independent effects, and alter host metabolic pathways, gene expression profiles, and immune cell function (15-19). Specifically, antibiotics can alter the host interferon response (19), alter immune cell homing to intestinal tissues (20, 21), and can have noncanonical effects on viruses, such as altering plaque size or exacerbating certain viral infections (15, 22). Intriguingly, it has been known for over 40 years that certain antibiotics have antiviral properties (23). Recently, Gopinath et al. demonstrated that aminoglycosides confer antiviral resistance *in vivo* to a variety of DNA and RNA viruses by altering the host immune response (19). Other antibiotics that have antiviral properties include macrolides (enterovirus A71) (24) and tetracyclines (respiratory syncytial virus) (25).

In this study, we examined the effects of antibiotics on systemic CVB3 and poliovirus infection in mice. We determined that treatment with a group of frequently used antibiotics for microbiota depletion studies (6) is capable of reducing systemic CVB3 and poliovirus titers following IP inoculation. Additionally, antibiotics altered early dissemination kinetics of CVB3, as well as late infection at 5 days post inoculation. Furthermore, we demonstrate that antibiotics reduced systemic CVB3 titers in microbiologically-sterile germ-free mice, suggesting a microbiota-independent effect.

## Results

### Antibiotic treatment reduces systemic enteric virus infection

Our previous work indicated that replication of orally-inoculated CVB3 was sensitive to antibiotics, suggesting that antibiotics have antiviral effects for virus within the gastrointestinal tract. To determine if antibiotics have antiviral activity for systemically delivered virus, we inoculated mice with CVB3 or poliovirus by IP injection. Because we sought to directly compare these two viral systems, we used mice transgenic for the human poliovirus receptor (PVR), which is required for poliovirus replication in mice. Mice with or without a weeklong administration of a combination of ampicillin, neomycin, metronadizole, vancomycin, and streptomycin were IP inoculated with 1×10^4^ PFU of coxsackievirus B3 Nancy strain (CVB3). Tissues were harvested at 72h post inoculation, and viral titers were quantified by plaque assay. We found that immunocompetent C57BL/6-PVR^+/+^ mice treated with antibiotics had significantly reduced CVB3 titers in several tissues, when compared to conventional mice, at 72h post inoculation (Fig. 1A). We next wanted to examine if this effect was dependent on type I interferon signaling by using mice that lack expression of the interferon alpha/beta receptor (IFNAR). We found that when immune-deficient C57BL/6-PVR^+/+^IFNAR^-/-^ mice were IP inoculated with CVB3, antibiotic treated mice had significantly reduced titers, when compared to conventional mice, at 72h post inoculation (Fig. 1B). These data suggest that antibiotics reduce systemic CVB3 titers in both immunocompetent and immunodeficient deficient mice.

**Figure 1.**
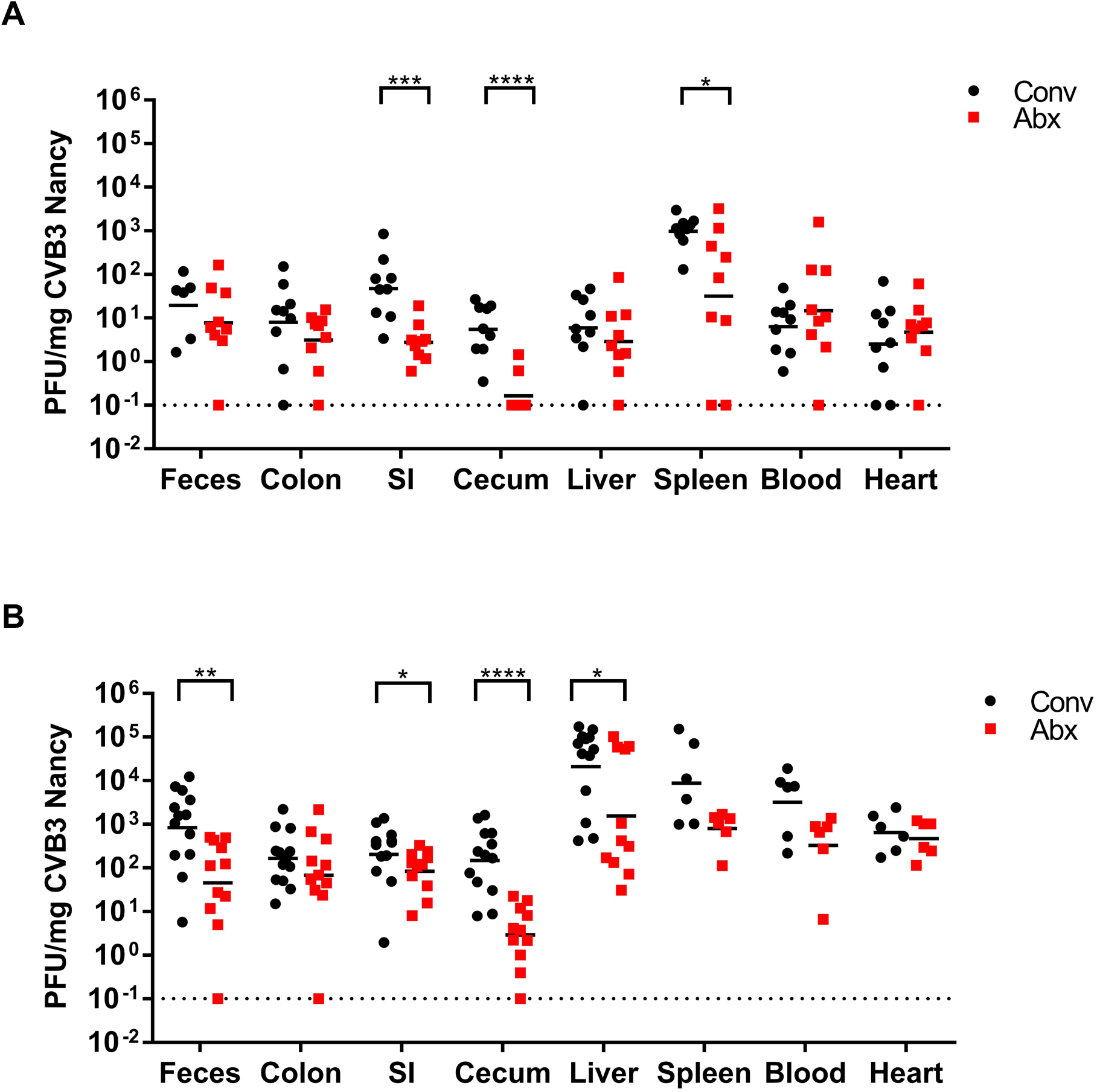
Effects of antibiotics on systemic CVB3 infection. Male C57BL/6 PVR^+/+^ IFNAR^+/+^ (A) or C57BL/6 PVR^+/+^ IFNAR^-/-^ (B) mice were treated with or without a combination of 5 antibiotics (ampicillin, neomycin, streptomycin, metronidazole, and vancomycin) for 5 days by oral gavage followed by *ad libitum* administration in drinking water for the duration of the experiment. Mice were IP inoculated with 1×10^4^ PFU CVB3 3-4 days post antibiotic gavage regimen. Tissues were collected at 72 hours post-inoculation and viral titers were determined by plaque assay. Dashed line represent limit of detection. Error bars represent geometric mean. *, *P*<0.05; **, *P*<0.001; ***, *P*<0.0001; ****, *P*<0.0001 (Mann-Whitney Test). n=6-13 mice per group. Data are from 3 independent experiments. SI, small intestine; Abx, antibiotics.

We next determined whether antibiotic treatment impacts systemic infection with a closely related enteric virus, poliovirus. We IP inoculated conventional or antibiotic-treated C57BL/6-PVR^+/+^IFNAR^-/-^ mice with 1×10^6^ PFU poliovirus and harvested tissues 72h post inoculation. We found that antibiotic-treated mice had significantly reduced poliovirus titers at 72h post inoculation, when compared to conventional mice (Fig. 2). Overall, these data indicate that antibiotic treatment reduces systemic poliovirus and CVB3 titers.

**Figure 2.**
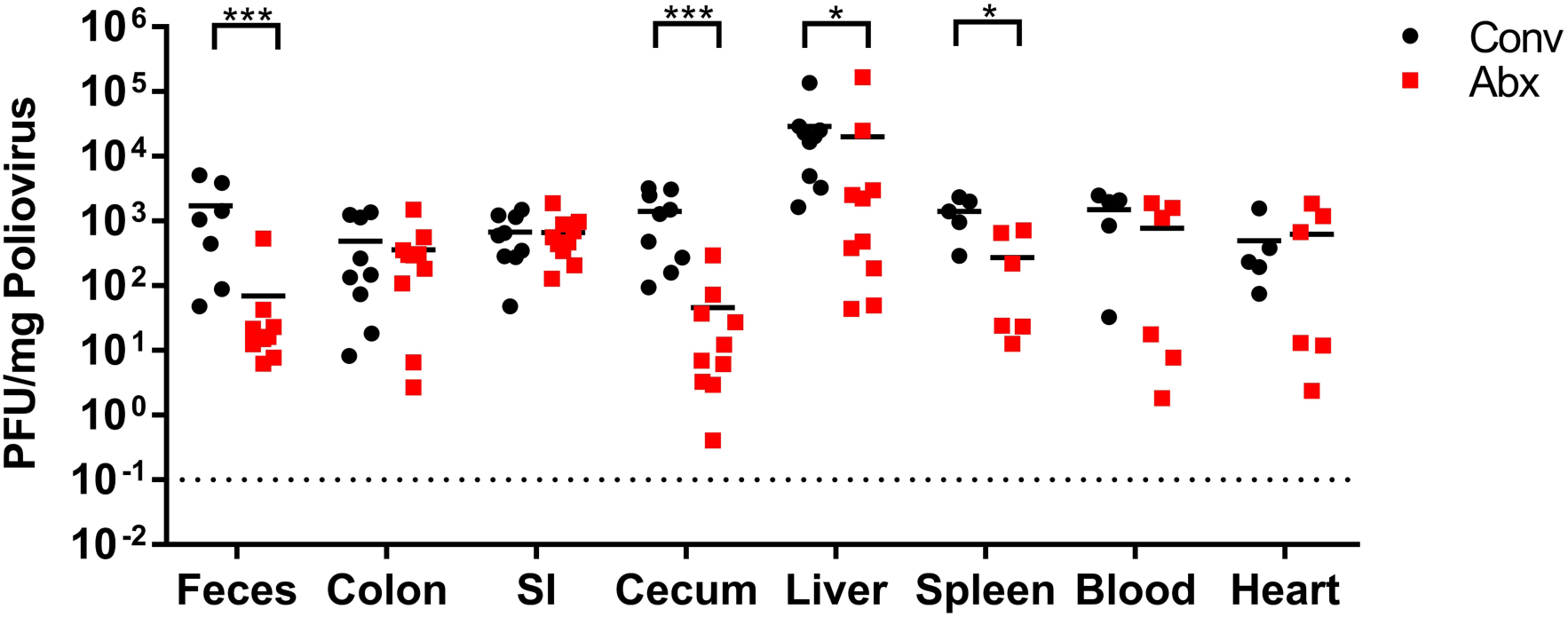
Effects of antibiotics on systemic poliovirus infection. Male C57BL/6 PVR^+/+^ IFNAR^-/-^ mice were treated with or without a combination of 5 antibiotics (ampicillin, neomycin, streptomycin, metronidazole, and vancomycin) for 5 days by oral gavage followed by *ad libitum* administration in drinking water for the duration of the experiment. Mice were IP inoculated with 1×10^6^ PFU poliovirus 3-4 days post antibiotic gavage regimen. Tissues were collected at 72 hours post-inoculation and viral titers were determined by plaque assay. Dashed line represent limit of detection. Error bars represent geometric mean. *, *P*<0.05; ***, *P*<0.0001 (Mann-Whitney Test). n=5-9 mice per group. Data are from 3 independent experiments. SI, small intestine; Abx, antibiotics.

### Antibiotics alter systemic CVB3 infection kinetics

Since antibiotic treatment was capable of reducing systemic CVB3 titers in mice at 72h post inoculation, we next investigated whether antibiotics alter CVB3 infection kinetics at earlier and later timepoints. We IP inoculated conventional or antibiotic-treated C57BL/6-PVR^+/+^IFNAR^-/-^ mice with 1×10^4^ PFU of CVB3 and harvested tissues at 24h, 72h, or 120h post inoculation and quantified the amount of virus by plaque assay. At 24h post inoculation we found that antibiotic-treated mice only had reduced titers in the feces (Fig. 3A-B). However, at 120h post inoculation, a majority of the tissues from antibiotic-treated mice had a significant decrease in viral titers (Fig. 3A-B). To further characterize the influence of antibiotics on the kinetics of systemic CVB3 infection, we examined whether antibiotics alter early CVB3 dissemination, prior to viral replication. We IP inoculated conventional or antibiotic-treated C57BL/6-PVR^+/+^IFNAR^-/-^ mice with 1×10^8^ PFU of CVB3 and harvested tissues at 1h post inoculation. We observed reduced titers in the majority of the tissues in antibiotic-treated mice at 1h post inoculation (Fig. 4), suggesting that early viral dissemination and/or clearance is altered by antibiotic treatment. Overall, these data indicate that antibiotics alter both CVB3 infection and dissemination kinetics.

**Figure 3.**
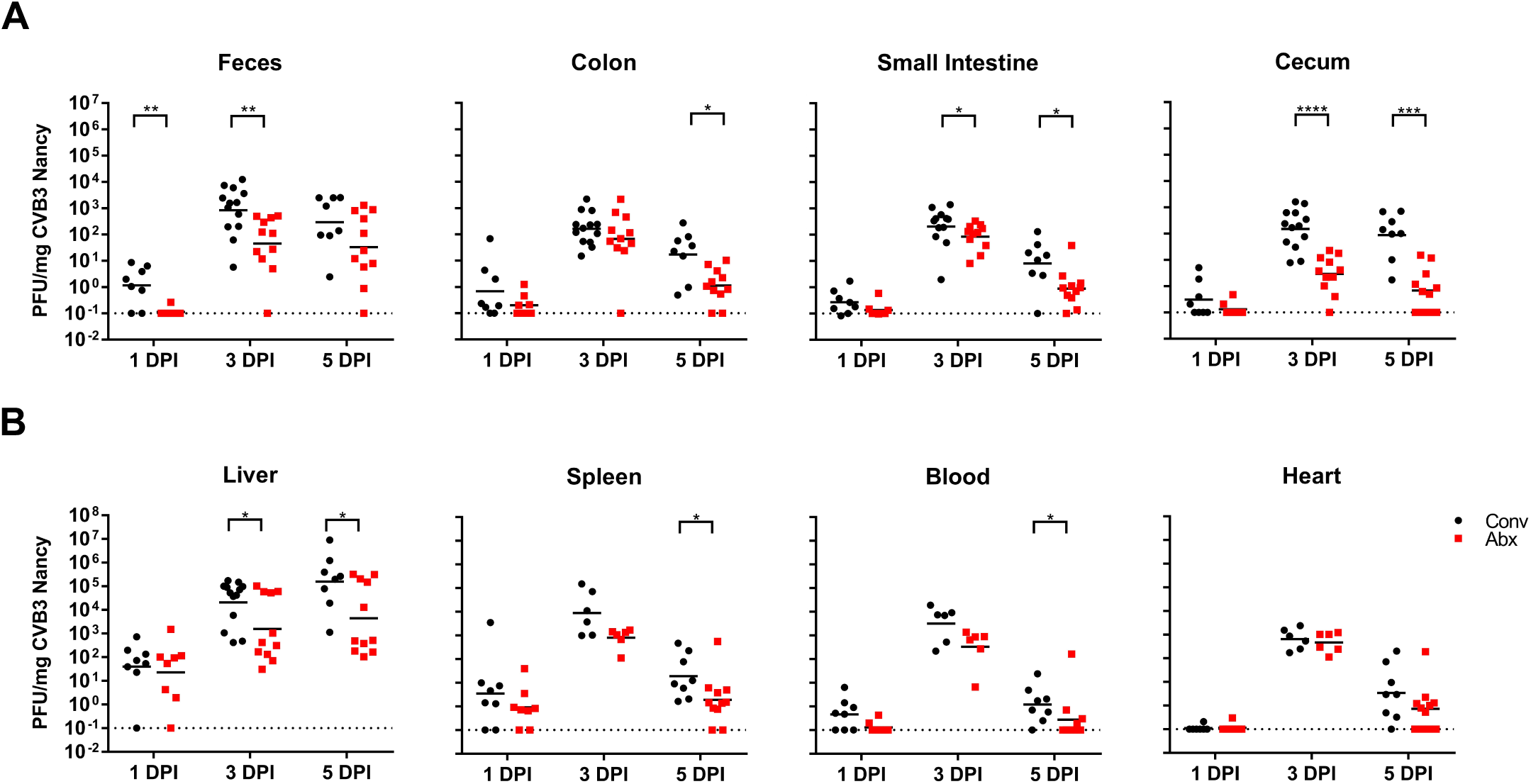
Antibiotics alter systemic CVB3 infection kinetics. Male C57BL/6 PVR^+/+^ IFNAR^-/-^ mice were treated with or without a combination of 5 antibiotics (ampicillin, neomycin, streptomycin, metronidazole, and vancomycin) for 5 days by oral gavage followed by *ad libitum* administration in drinking water for the duration of the experiment. Mice were IP inoculated with 1×10^4^ PFU CVB3 3-4 days post antibiotic gavage regimen. Tissues were collected at 1, 3, or 5 days post-inoculation and viral titers were determined by plaque assay. Dashed line represent limit of detection. Error bars represent geometric mean. *, *P*<0.05; **, *P*<0.001; ***, *P*<0.0001; ****, *P*<0.0001 (Mann-Whitney Test). n=6-13 mice per group. Data are from 3 independent experiments. SI, small intestine; Abx, antibiotics.

**Figure 4.**
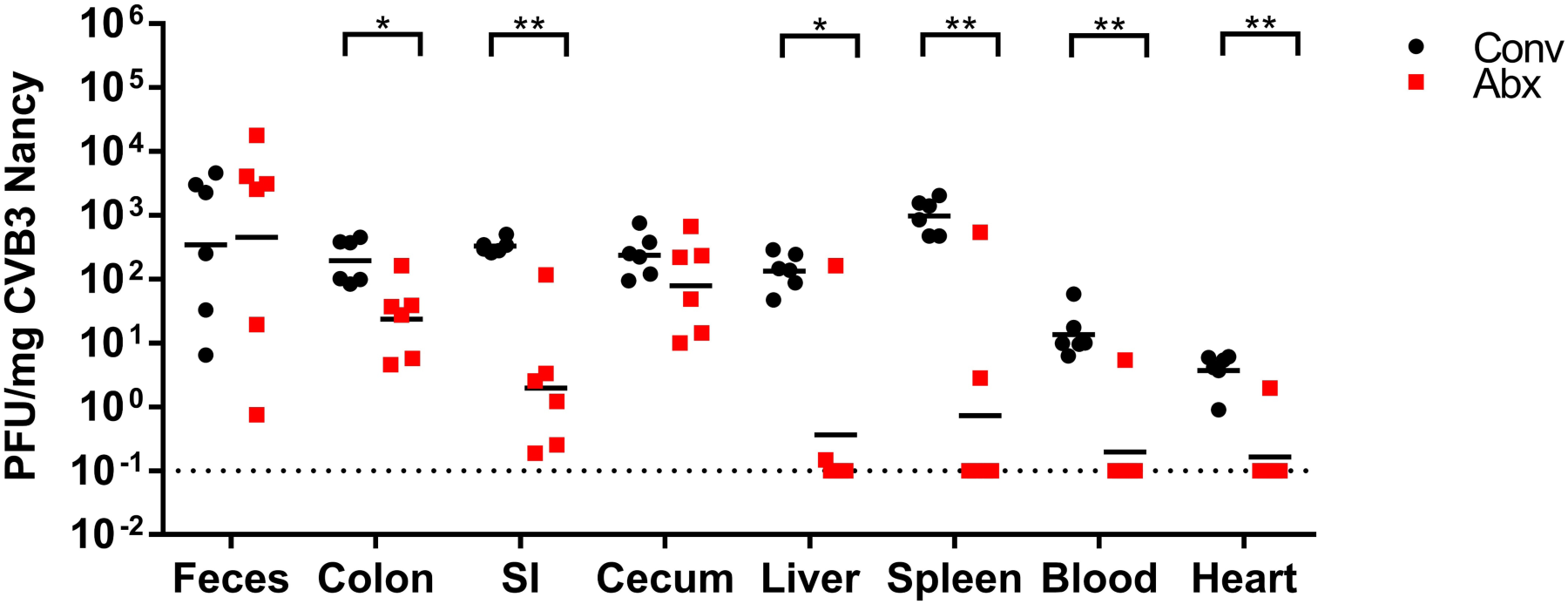
Antibiotics alter early CVB3 dissemination kinetics. Male C57BL/6 PVR^+/+^ IFNAR^-/-^ mice were treated with or without a combination of 5 antibiotics (ampicillin, neomycin, streptomycin, metronidazole, and vancomycin) for 5 days by oral gavage followed by *ad libitum* administration in drinking water for the duration of the experiment. Mice were IP inoculated with 1×10^8^ PFU CVB3 3-4 days post antibiotic gavage regimen. Tissues were collected at 60 minutes post-inoculation and viral titers were determined by plaque assay. Dashed line represent limit of detection. Error bars represent geometric mean. *, *P*<0.05; **, *P*<0.001 (Mann-Whitney Test). n=6 mice per group. Data are from 2 independent experiments. SI, small intestine; Abx, antibiotics.

### Treatment with individual antibiotics reduces CVB3 titers in the cecum

Given that the cocktail of five antibiotics was capable of reducing CVB3 titers, we next wanted to determine which of the individual antibiotics was responsible for the antiviral effects. To test this, we IP inoculated conventional or individual antibiotic-treated C57BL/6-PVR^+/+^IFNAR^-/-^ mice with 1×10^4^ PFU of CVB3 and harvested the ceca at 72h post inoculation. We chose to focus on the ceca since we previously observed decreased cecal titers at 72h- and 120h- post inoculation with CVB3 and at 72h post inoculation with poliovirus. We found that each of the five individual antibiotics could reduce CVB3 cecal titers to varying degrees (Fig. 5). Overall, these data indicate that each individual antibiotic in the cocktail is capable of reducing CVB3 cecal titers.

**Figure 5.**
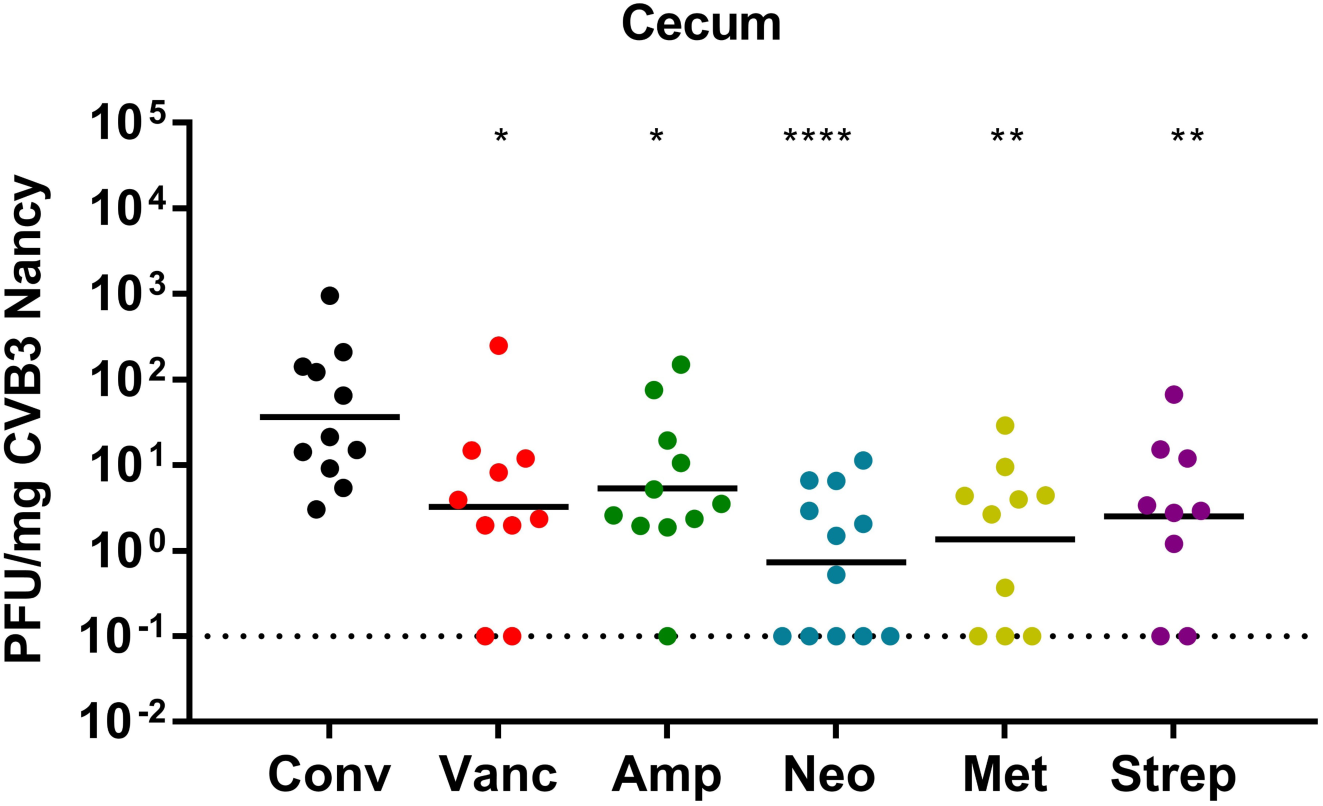
Treatment with individual antibiotics reduce CVB3 cecal titers. Male C57BL/6 PVR^+/+^ IFNAR^-/-^ mice were treated with or without each of the 5 antibiotics (ampicillin, neomycin, streptomycin, metronidazole, and vancomycin) individually for 5 days by oral gavage followed by *ad libitum* administration in drinking water for the duration of the experiment. Mice were IP inoculated with 1×10^4^ PFU CVB3 3-4 days post antibiotic gavage regimen. Ceca were collected at 72 hours post-inoculation and viral titers were determined by plaque assay. Dashed line represent limit of detection. Error bars represent geometric mean. *, *P*<0.05; **, *P*<0.001; ****, *P*<0.0001 (Mann-Whitney Test). n=9-11 mice per group. Data are from 3 independent experiments. Conv, conventional; Vanc, vancomycin; Amp, ampicillin; Neo, neomycin; Met, metronadizole; Strep; streptomycin.

### Antibiotics reduce CVB3 titers in germ-free mice

Given that the antibiotic cocktail was able to reduce CVB3 titers in conventionally raised mice upon systemic infection, we next sought to determine if antibiotics could also reduce CVB3 titers in a microbiota-independent fashion. To test this, microbiologically-sterile germ-free C57BL/6 mice were treated with or without a weeklong administration of a combination of ampicillin, neomycin, metronadizole, vancomycin, and streptomycin and were then intraperitonially inoculated with 1×10^4^ PFU of CVB3. Tissues were harvested at 72h post inoculation, and viral titers were quantified by plaque assay. We observed that germ-free mice treated with antibiotics had significantly reduced CVB3 titers in several tissues, when compared to mock-treated mice, at 72h post inoculation (Fig. 6). These data suggest that antibiotics reduce CVB3 titers in a microbiota-independent manner.

**Figure 6.**
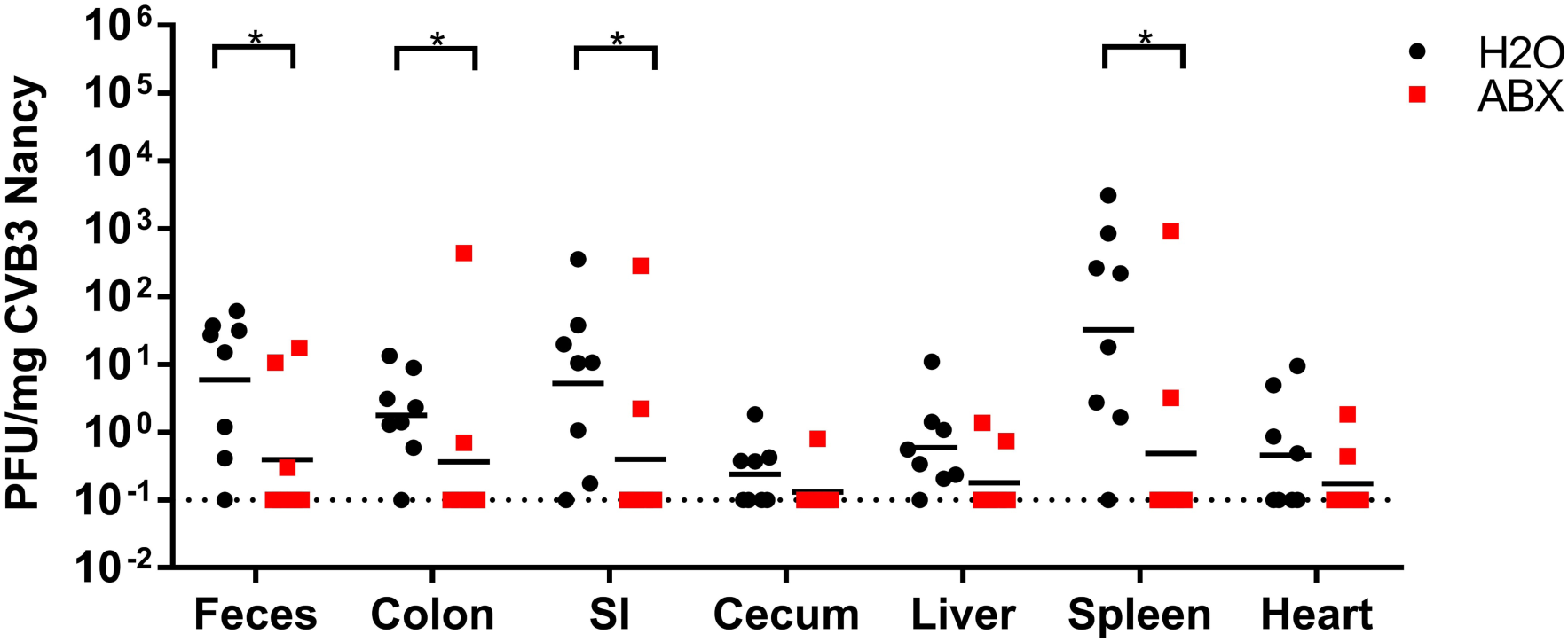
Antibiotics reduce systemic CVB3 titers in germ-free mice. Male C57BL/6 germ-free mice were treated with or without a combination of 5 antibiotics (ampicillin, neomycin, streptomycin, metronidazole, and vancomycin) for 5 days by oral gavage followed by *ad libitum* administration in drinking water for the duration of the experiment. Mice were IP inoculated with 1×10^4^ PFU CVB3 3-4 days post antibiotic gavage. Tissues were collected at 72 hours post-inoculation and viral titers were determined by plaque assay. Dashed line represent limit of detection. Error bars represent geometric mean. *, *P*<0.05 (Mann-Whitney Test). n=8 mice per group. Data are from 2 independent experiments. SI, small intestine; Abx, antibiotics.

## Discussion

Work performed in several labs has shown that antibiotic treatment reduces replication and pathogenesis of orally inoculated enteric viruses (6, 11-14). However, the influence of antibiotics on systemic enteric virus infection is unclear. Similarly, it is not known whether microbiota-independent effects of antibiotics influence enteric virus infection. To bypass direct interactions with enteric viruses and bacteria that occur following oral inoculation, mice were IP injected with CVB3 or poliovirus. In this study we show that a tissue-specific antiviral effect is observed when conventionally-raised or microbiologically-sterile mice are treated with antibiotics and IP infected with CVB3.

There has been a growing body of evidence that antibiotics can have a wide range of effects on the host (26). One study of particular note determined that aminoglycoside antibiotics are capable of altering the innate immune response in mice, which resulted in antiviral activity against both DNA and RNA viruses (19). They also found that these antibiotics function in a microbiota-independent manner (19), similar to our observations. Interestingly, two out of the five antibiotics in our cocktail are aminoglycoside antibiotics, both of which individually reduced systemic CVB3 cecal titers. Administration of aminoglycoside antibiotics can immune cell homeostasis in various tissues (19-21). Of note, a single dose of the aminoglycoside antibiotic streptomycin promoted the infiltration of neutrophils and inflammatory monocytes to the cecum (20). Interestingly, oral CVB3 infection is highly sensitive to antibiotics, where a single dose of streptomycin can significantly decrease viral replication and pathogenesis (6). Given that we and others have shown noncanonical effects of antibiotics, it is possible that systemic enteric virus infection could be influenced by antibiotic-induced innate immune response and/or immune cell infiltrates, although future experiments will be needed to test this hypothesis.

In conclusion, we found that antibiotics alter systemic enteric viral infection in both immunocompetent and type I interferon deficient mice. Antibiotics were also capable of reducing systemic CVB3 titers in microbiologically-sterile germ-free mice, suggesting a microbiota-independent effect. These findings imply that standard antibiotic treatment regimens for bacterial infections could also impact the course of enteric virus infections.

## MATERIALS AND METHODS

### Viruses and Cells

HeLa cells were grown in Dulbecco’s modified Eagle’s medium supplemented with 10% calf serum (Sigma-Aldrich) and 1% penicillin-streptomycin (Sigma-Aldrich). The CVB3-Nancy infectious clone was obtained from Marco Vignuzzi (Pasteur Institute, Paris, France). The poliovirus infectious clone was serotype 1 Mahoney (27). Viral stocks were prepared as previously described (28). Viral titers were determined by plaque assay using HeLa cells as previous described (6).

### Mice, treatments, and inoculations

All animals were handled according to the *Guide for the Care and Use of Laboratory Animals* endorsed by the National Institutes of Health and in accordance with University of Texas Southwestern Medical Center IACUC-approved protocols. C57BL/6-PVR^+/+^-IFNAR^+/+^ and C57BL/6-PVR^+/+^-IFNAR^−/−^ mice were obtained from S. Koike (Tokyo, Japan). Note that all mice used in this study, with the exception of germ-free C57BL/6 mice, express the human poliovirus receptor as a transgene. Mice were maintained in specific pathogen-free conditions. Throughout, cage randomization of different litters was used to limit differences caused by litter-to-litter and microbiota variability. For example, the mice from each cage were split into two groups—one treated with antibiotics and one not treated with antibiotics. Microbiologically-sterile germ-free C57BL/6 mice were maintained in gnotobiotic chambers. For CVB3 infections, germ-free mice were housed in the BSL2+ facility in sterile cages with autoclaved food, water, and bedding for the week of the experiment. 8-12 week old male C57BL/6-PVR^+/+^-IFNAR^+/+^ or C57BL/6-PVR^+/+^-IFNAR^−/−^ mice were orally administered a combination of five antibiotics: vancomycin, ampicillin, neomycin, metronidazole, and streptomycin via oral gavage for 5 days (10 mg of each antibiotic per mouse per day) followed by *ad libitum* administration in drinking water (vancomycin: 500 mg/L; ampicillin, neomycin, metronidazole, and streptomycin: 1 g/L;) for the duration of the experiment. Control mice were treated with oral gavage of H_2_O to mimic antibiotic treatment. For IP inoculation, 8- to 12-week-old mice were inoculated with 1 × 10^4^ PFU of CVB3 or 1 × 10^6^ PFU of poliovirus in 150µL of sterile PBS. For dissemination kinetics, 8- to 12-week-old mice were IP injected with 1 × 10^8^ PFU of CVB3. Upon disease onset mice were euthanized by isoflurane followed by cervical dislocation. Male mice were used throughout, since CVB3 has a sex-dependent enhancement of replication and lethality (7).

### Sample processing and titer analysis

Tissues were aseptically removed from mice. Samples were homogenized in PBS using 0.9- to 2.0-mm stainless steel beads in a Bullet Blender (Next Advance) and freeze-thawed 3 times in liquid nitrogen to release intracellular virus. Cellular debris was removed by centrifugation at 13,000 RPM for 3 minutes. The amount of virus present in supernatant was quantified by plaque assay and normalized to the weight of each tissue (6).

### Statistical Analysis

The difference between groups were examined by the unpaired two-tailed Student *t* test. Error bars in figures represent the geometric mean. A *P* value of <0.05 was considered significant. All analyses of data were performed using GraphPad Prism version 7.00 for Windows, GraphPad Software, La Jolla, CA.

## Funding Information and Acknowledgements

Work in J.K.P.’s lab is funded through NIH NIAID grant R01 AI74668, a Burroughs Wellcome Fund Investigators in the Pathogenesis of Infectious Diseases Award, and a Faculty Scholar grant from the Howard Hughes Medical Institute. MWA was supported in part by NIH NIAID grant T32 AI007520.

